# Alteration of Gut Microbiome in Lung Cancer Patients

**DOI:** 10.1101/640359

**Authors:** Li Ming, Yu Fang, Chen Xiaohui, Zhou Huan, Wei Xiaoqing, Liu Yinhui, Liu Yuanyu, Tang Li, Yuan Jieli, Wen Shu, Chen Jun

## Abstract

Lung cancer is the leading cause of cancer death. Better understanding of factors and pathways involved in lung cancer is needed to improve diagnose and treatment strategies. Recent studies have provided insights into the possible correlation between intestinal dysbiosis and cancer development. Although the immunological relationship between gut and lung had been suggested by many researches, however, to date, no study had investigated the characterization of gut microbiome in treatment naïve lung cancer patients, whether it is distinct from that of health individuals and contribute to the onset and development of lung cancer remain unclear. In this study, we investigated whether gut microbiome of lung cancer patients (LC, n=28) is altered compare with that of matched healthy individuals (HC, n=19) by high throughout sequencing of the V3-V4 regions of 16S rDNA in their fecal samples. We also identified microbiota signatures specific for different histological types of lung cancer, including SSC, ADC, and SCLC. The gut microbiome of lung cancer patients is characterized by decreased relative abundance of *Prevotella*, and increased bacteria groups such as *Actinomyces*, and *Streptococcus*, etc. We also detected a mild structural shift in gut microbiome between ADC and SCLC patients. Our results showed that the gut microbiome of lung cancer patients altered significantly compared with healthy individuals. However, the association between microbial dysbiosis and lung cancer is not clearly understood, future studies involving larger cohorts and metagenomics, or metabolomics, may elucidate the correlations between gut microbiota and lung cancer development.

**IMPORTANCE:** This is the first report to show the alteration of gut microbiome in lung cancer patients. Our results showed that the gut microbiome of lung cancer patients altered significantly compared with healthy individuals.

## INTRODUCTION

Lung cancer is the leading cause of cancer death (1). There are estimated 1.82 million new cases of lung cancer globally, which constitutes nearly 13% of all newly diagnosed cancer cases annually (2). More than one-third of lung cancer worldwide occurring in China, where 733,000 new cases of lung cancer are diagnosed, and about 591,000 Chinese people died from it each year (3-4). Better understanding of factors and pathways involved in lung cancer is urgently needed to improve treatment strategies.

As a heterogeneous disease, many factors are involved in the onset and development of lung cancer. Although smoking is considered as an important factor, only 10–15% of smokers develop cancer (5), which highlights other influences, such as the involvement of microbial communities. Recent studies showed that changes in the lung microbiome may be relevant for progression and exacerbations in lung cancer (6,7). By comparison of microbiome in bronchoalveolar lavage fluid of patients with lung cancer with benign mass like lesions, Lee *et al.* found that the genera *Veillonella* and *Megasphaera* are more abundant in lung cancer patients, which may serve as potential biomarkers for the disease detection/classification (8). A recent research, in which surgical lung tissue samples were used, concluded that the microbiota of the lung cancer is unique, with the genus *Thermus* more abundant in tissue from advanced stage (IIIB, IV) patients, while *Legionella* is higher in patients who develop metastases (9). These studies provide insights into the possible correlation between microbiota and lung cancer development.

However, lung microbiota doesn’t seem to be the only microbial factor contributing to the development of lung cancer. The influence of gut microbiota on lung immunity has been vastly explored and several studies have linked changes in the gut microbiome with lung diseases (10,11). For example, gut flora is responsible for inducing lung inflammatory reaction against bacterial challenge and enhancing neutrophils infiltration through TLR4 in mice (12,13). Antibiotic treatment can cause overgrowth of particular fungal species in the gut and promote allergic airway inflammation via fungi-induced prostaglandin E2 (14). Conversely, the lung microbiota also influences the gut microbiota through the blood stream. Acute lung injury (ALI) can disrupt the lung microbiota, induces a transient translocation of bacteria into blood and causes an acute increase of bacterial load in cecum (15). These studies emphasized the important role of gut-lung axis in development of diseases (16). However, to date, no study had investigated the characterization of gut microbiome in treatment naïve lung cancer patients, whether it is distinct from that of health individuals and contribute to the onset and development of lung cancer remain unclear.

In this study, we investigated whether gut microbiome of lung cancer patients (LC, n=28) is altered compare with that of matched healthy individuals (HC, n=19) by high throughout sequencing of the V3-V4 regions of 16S rDNA in their fecal samples. We also identified microbiota signatures specific for different histological types of lung cancer, including SSC, ADC, and SCLC.

## RESULTS

### Characteristics of participants

A total of 50 participants enrolled in the Department of Medical Oncology, The Second Affiliated Hospital of Dalian Medical University (Dalian, China). Participants were excluded after fecal sample collection because of antibiotic use, received chemotherapy, or combined with other diseases such as diabetes mellitus (Fig. 1). Finally, a total of 28 LC patients were remained, they were mainly males (83%) with a median age of 65 years old (Table 1) and with an average smoking index of 606.79±127.1. 19 healthy controls (HC) were included for age and gender matching, with an average smoking index of 350.00±72.1 (P > 0.05 compared with LC group) (details in Table 1). More than 64% of the enrolled LC patients were smokers, with a high smoking index about 606.79±127.1. Among them, 28.57% (8/28) of the patients were diagnosed as SCLC, and 71.43% (20/28) were NSCLC. Among the NSCLC patients, 21.43% (6/28) were diagnosed as squamous cell carcinoma and 50.00% (14/28) were adenocarcinoma.

**TABLE 1.**
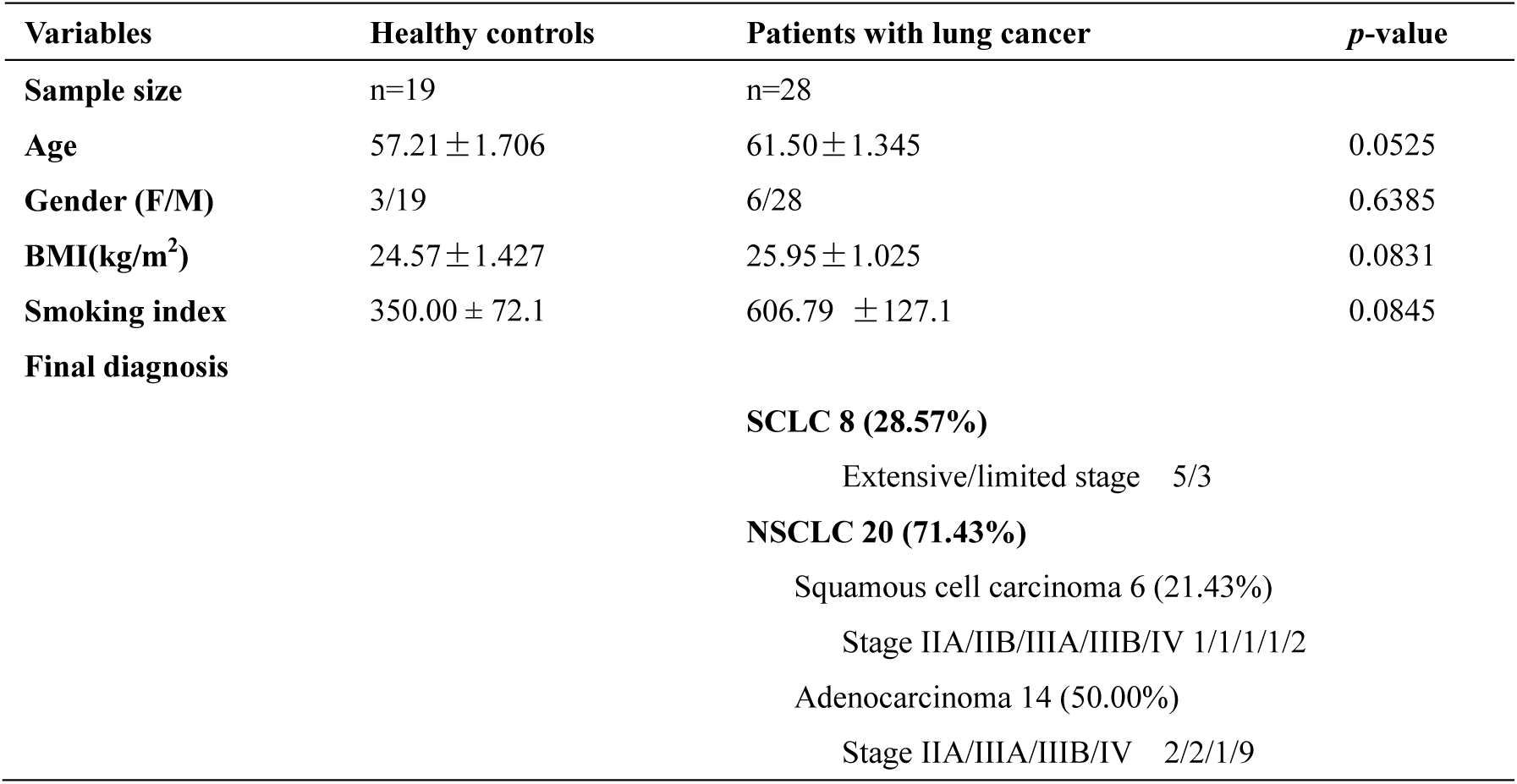
Characteristics of the study population

| Variables | Healthy controls | Patients with lung cancer | *p*-value |
| --- | --- | --- | --- |
| Sample size | n=19 | n=28 |  |
| Age | 57.21 ± 1.706 | 61.50 ± 1.345 | 0.0525 |
| Gender (F/M) | 3/19 | 6/28 | 0.6385 |
| BMI(kg/m<sup>2</sup>) | 24.57 ± 1.427 | 25.95 ± 1.025 | 0.0831 |
| Smoking index | 350.00 ± 72.1 | 606.79 ± 127.1 | 0.0845 |
| Final diagnosis |  |  |  |
|  |  | **SCLC 8 (28.57%)** |  |
|  |  | Extensive/limited stage 5/3 |  |
|  |  | **NSCLC 20 (71.43%)** |  |
|  |  | Squamous cell carcinoma 6 (21.43%) |  |
|  |  | Stage IIA/IIB/IIIA/IIIB/IV 1/1/1/1/2 |  |
|  |  | Adenocarcinoma 14 (50.00%) |  |
|  |  | Stage IIA/IIIA/IIIB/IV 2/2/1/9 |  |

**FIG 1.**
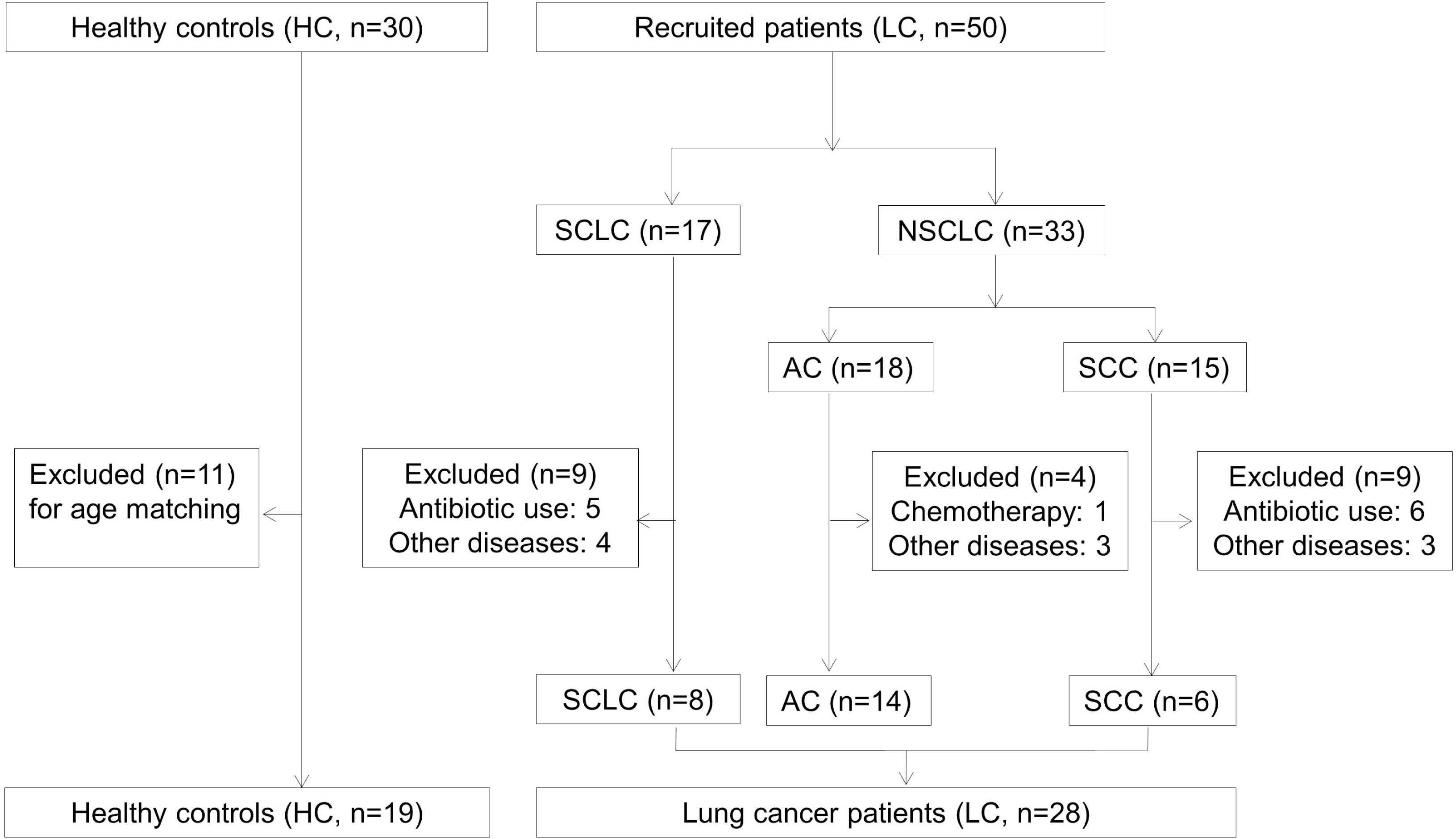
The recruitment of participants and the process of sample collection.

### The overall structure of gut microbiome in LC patients

By sequencing of the 16S ribosomal RNA gene, we found that The HC and LC groups share about 2368 same Operational Taxonomic Units (OTUs), but there are 372 OTUs were obtained specifically in the HC group, and 202 were obtained specifically in the LC group (Fig. 2A). We observed significant decrease in alpha diversity of gut microbiota in LC group, which expressed by the ACE and Chao1 index (Fig. 2B and C). Whereas the Shannon diversity index and the Simpson index did not show significant differences between two groups (Fig. S1). Changes in the relative abundance of gut microbes in LC patients were observed not only on the phylum level, but also on the levels of order, class, and family (Fig. S4). At genus level, significant decreased abundance of *Prevotella* and elevated abundance of *Bacteroides*, and *Ruminococcus* etc. were detected in LC patients (Fig. 2D).

**FIG 2.**
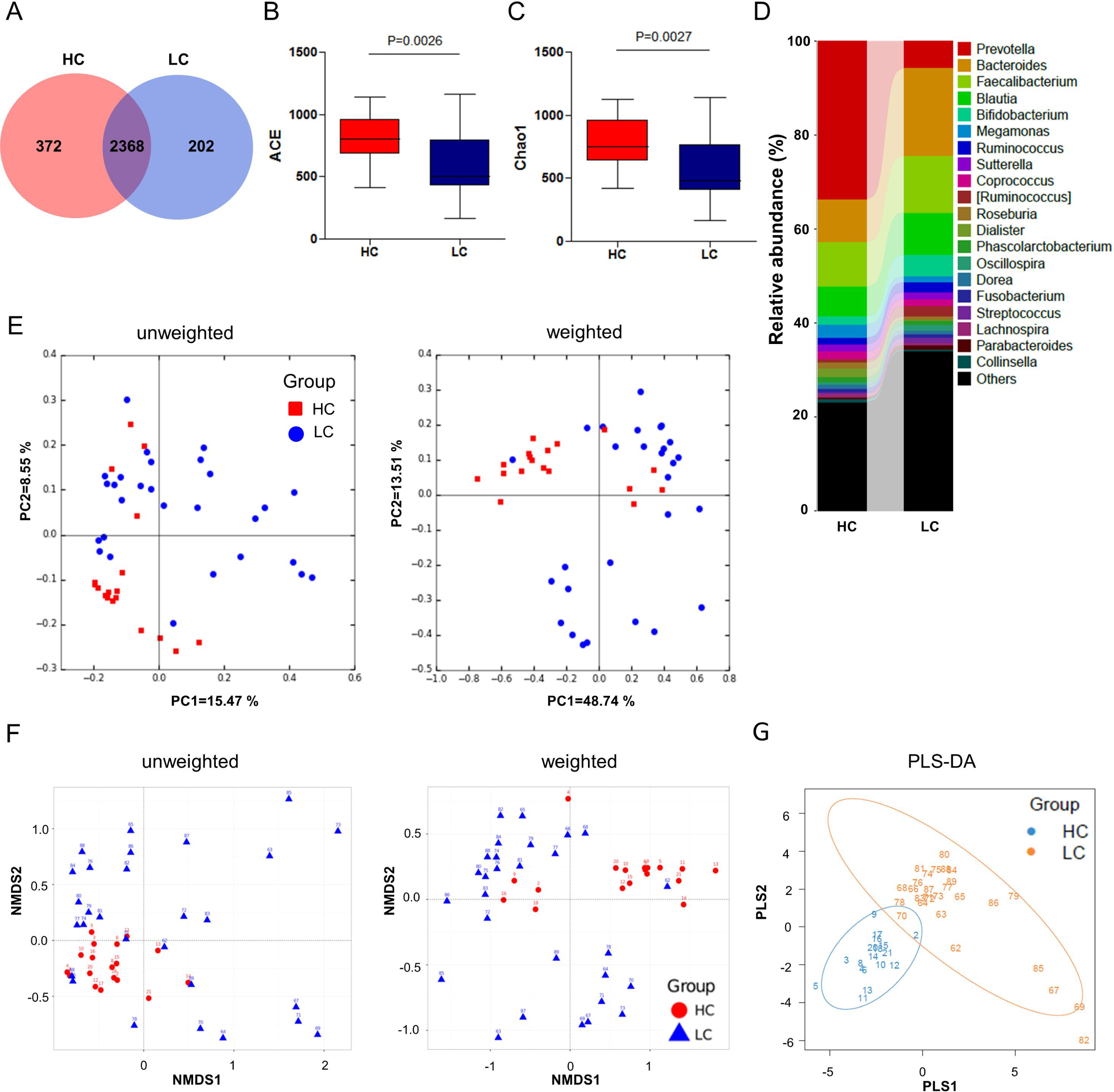
The intestinal bacterial composition in HCs and LC patients. **(A)** Venn diagram of shared and independent bacterial OTUs in HC and LC groups. **(B)** Comparison of the ACE and Chao1 index of HC and LC groups. **(C)** The bacterial composition in different groups at Genus level. **(D)** Principal Coordinate Analysis (PCoA) based on weighted Unifrac distances among different samples. **(E)** NMDS based on weighted Unifrac distances among different samples. **(F)** PLS-DA of the gut microbiome in HC and LC patients.

The beta diversity metrics from the control and LC individuals also showed strong grouping pattern. Although significant inter-individual variation exists among patients and the healthy controls, the fecal microbiota of the two groups still separated clearly according to community composition using Principal component analysis (PCA, Fig. S3), and unweighted/weighted UniFrac Principal coordinates analysis (PCoA) (Fig. 2E). These differences were also observed by the two-dimensional Nonmetric Multidimensional Scaling (NMDS) based on unweighted/weighted UniFrac (Fig. 2F). Most of the samples from each group clustered together as evaluated by Hierarchical clustering based on Weighted UniFrac by the method of Unweighted pair-group method with arithmetic means (UPGMA, Fig. S4). Especially, when analyzed by the method of Partial Least Squares Discriminant Analysis (PLSDA), we observed a significant separation between the LC patients and HCs (Fig. 2G).

### Altered microbiota composition in LC patients

The alteration of gut microbiome in LC patients was further proved by the LEfSe approach, which identified the key phylotypes responsible for the difference between the two groups. *Actinobacteria, Bacilli, Ruminococcus, Streptococcus*, and *Mycobacteriaceae*, etc, which were most abundant in the LC group, and *Prevotella, Bacteroidetes*, and *Dialister*, which were most abundant in the HCs, were the dominant phylotypes that contributed to the difference between the intestinal microbiota of LC patients and HCs (Fig. 3A and B). The significantly elevated relative abundance of *Actinomyceae, Streptococcus*, and *Ruminococcus*, and decreased abundance of *Prevotellaceae* were observed in the gut microbiome of most of the LC patients, which suggested a highly consistence among different individuals (Fig. S5).

**FIG 3.**
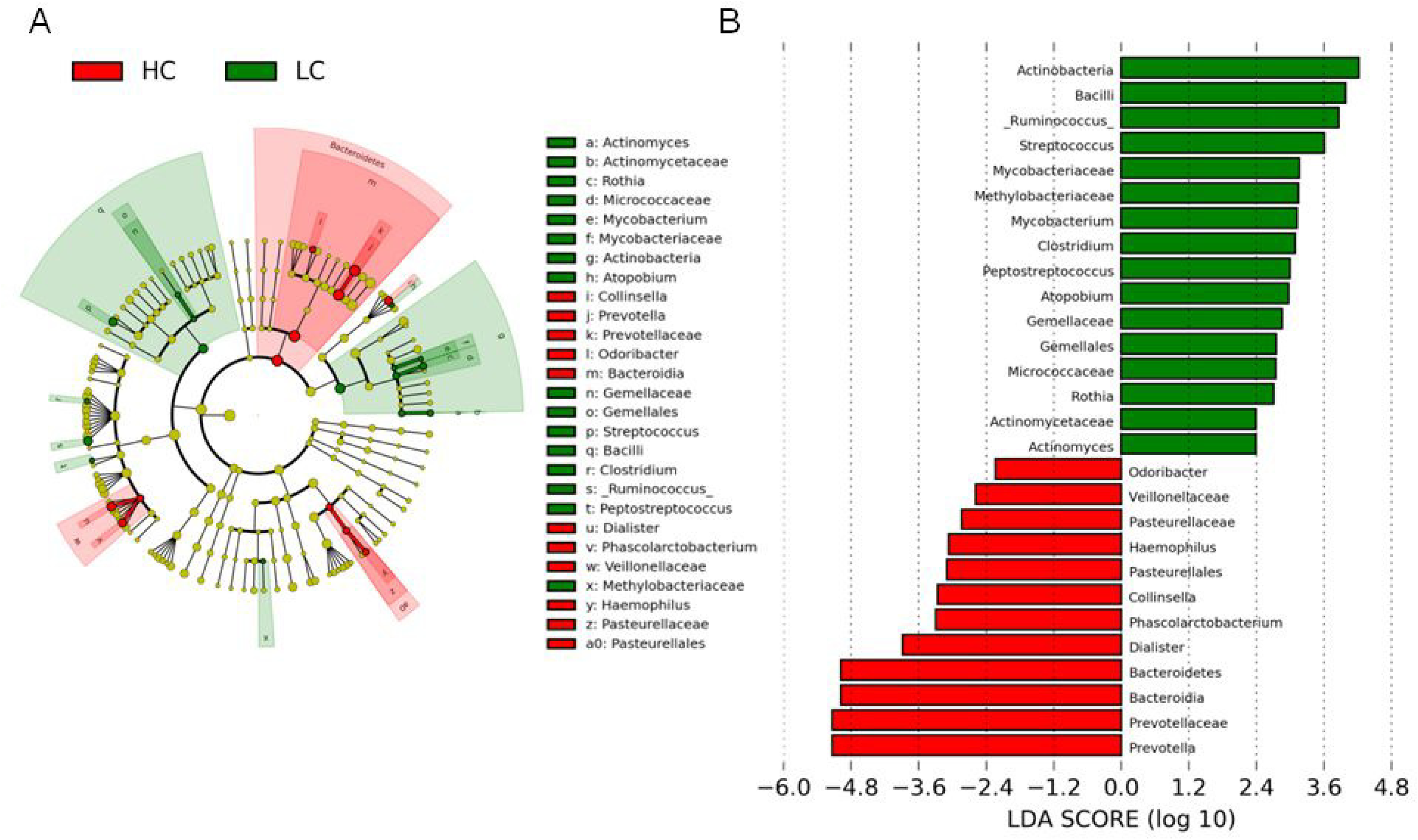
LEfSe analysis of gut microbiota in HC and LC groups. **(A)** LEfSe identified the most differentially abundant bacterial taxons among groups. Group-specific enriched taxa are indicated with a positive LDA score bar with different colors. Only taxa meeting an LDA significant threshold >2 are shown. **(B)** Taxonomic cladogram obtained from LEfSe analysis of 16S rDNA sequences. The brightness of each dot is proportional to its effect size.

Using the method of Metastats, we found a significant decrease in the abundance of *Bacteroidetes* (phylum), *Bacteroidia* (class), *Bacteroidales* (order) and elevated abundance of *Firmicutes* (phylum), *Bacilli* (class), *Actinomycetales* (order), *Bacillales* (order), *Lactobacillales* (order) in gut of the LC patients (Fig. S6). On the family level, significant elevation of the relative abundance of *Streptococcaceae, Actinomycetaceae*, decreased abundance of the *Prevotellaceae* and *Veillomellaceae* were observed in gut of LC patients compared with HCs (Fig. 4A). These changes may mainly due to the changing of genus such as the *Streptococcus, Actimomyces*, and *Prevotella* etc (Fig. 4B). In addition, we also detected elevation of genera such as *Ruminococcus, Rothia, Bacillus, Peptostreptococcus, Mycoacterium*, etc, and decreased abundance of *Dialister* in gut of LC patients, which are consistent with LEfSe analysis results.

**FIG 4.**
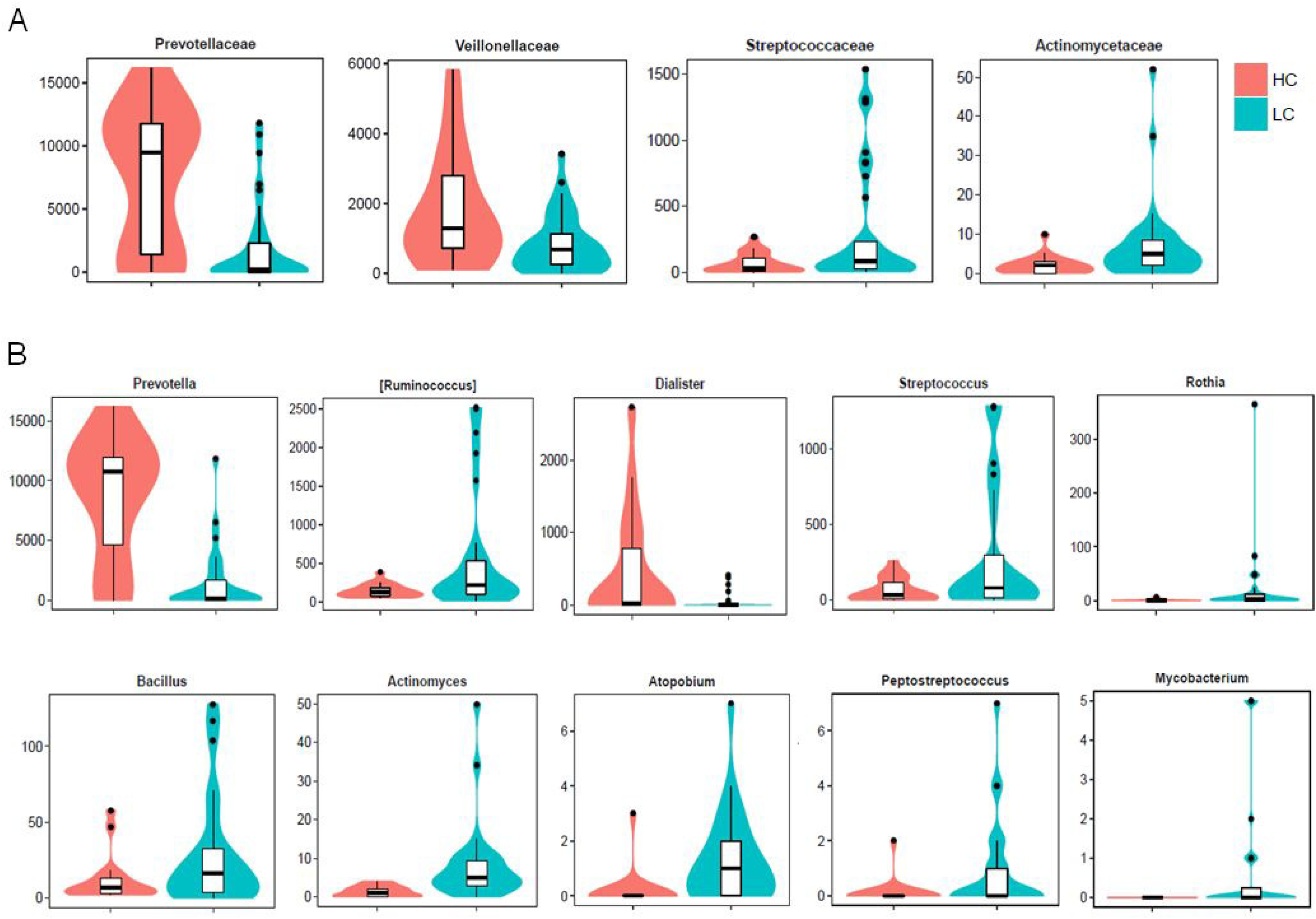
The specific bacterial groups that are significantly changed in LC patients compared with HCs detected by the MetaStat method. **(A)** Families, **(B)** Genera.

### Comparison of gut microbiome in LC patients with specific histological types

We further performed a detailed comparison of the gut microbiome in lung cancer patients according to different histological types, including adenocarcinoma (Group A, n=14), squamous cell carcinoma (Group B, n=6), and SCLC (Group C, n=8). Only the index of alpha diversity (ACE) in SCLC patients is significantly lower than control (Fig. 5A). Other groups showed no significant differences compared either with HC or other groups, although the average indexes of each of the specific histological type are obviously lower than that of the HCs (Fig. 5A and Fig. S7). The beta diversity analysis by PCoA and NMDS showed no obvious separation between groups (Fig. 5B and C). When analyzed by the method of PLSDA, we observed a mild separation between group A and C, which suggested that the gut microbiome of SCLC patients may differ from that of LC patients with adenocarcinoma (Fig. 5D). The taxonomy-based comparison at the genus level showed that, *Bifidobacterium, Clostridium*, and *Prevotella*, etc, are the dominant phylotypes in group A, but were significantly reduced in the other two groups. In group B, the dominant genera including *Ruminococcus, Lachnospira*, and *Lactobacillus*, etc, which are less abundant in group A and C. in addition, the genera of *Streptococcus, Anaerotruncus*, and *Bacillus*, etc are more abundant in group C when compared with group A and B (Fig. S8).

**FIG 5.**
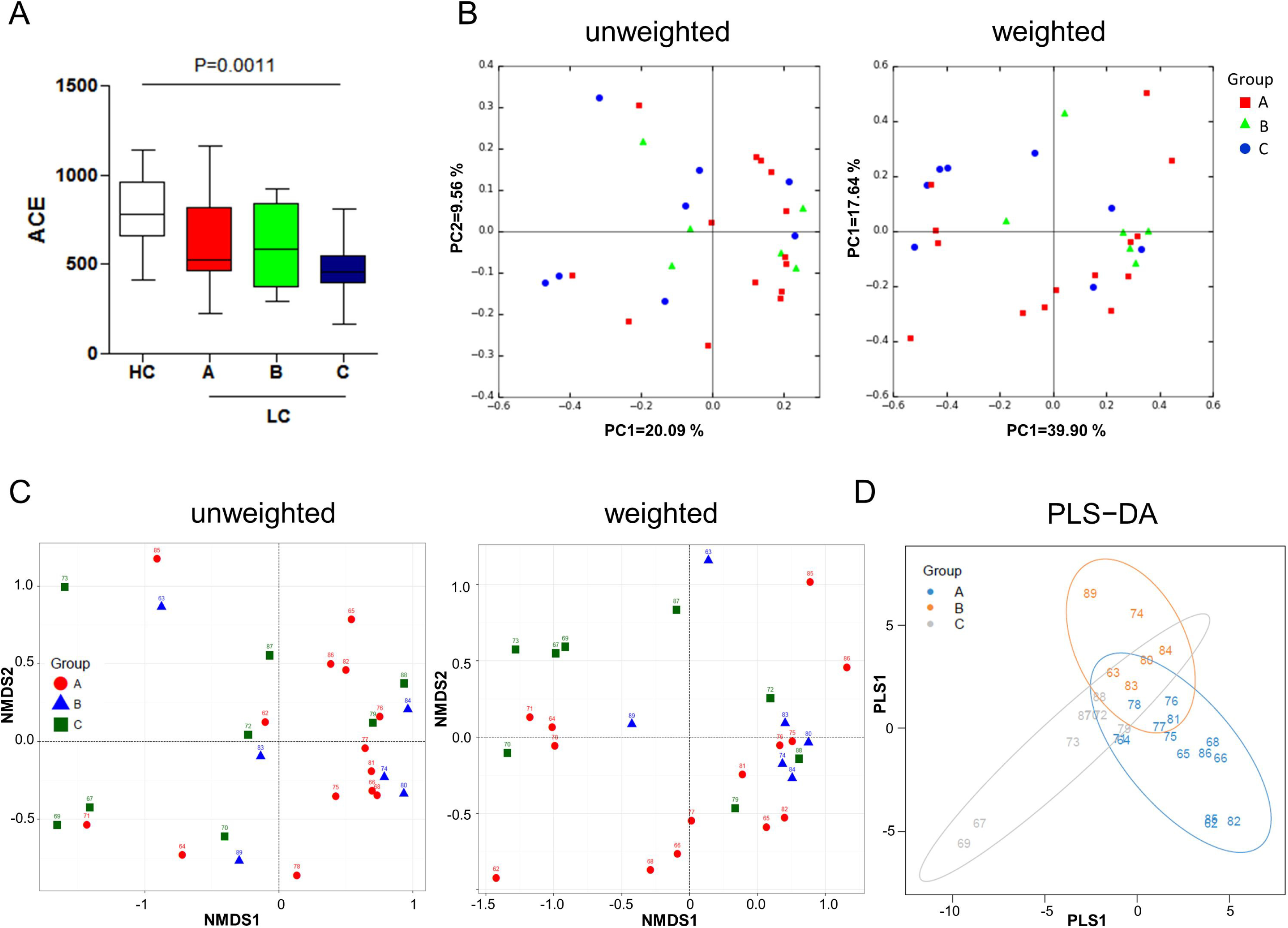
Comparison of gut microbiome in LC patients with specific histological types. **(A)** Comparison of the ACE index of different groups. **(B)** PCoA based on unweighted and weighted Unifrac distances among different samples. **(C)** NMDS based on unweighted and weighted Unifrac distances among different samples. **(D)** PLS-DA of the gut microbiome in LC patients with specific histological types.

## DISCUSSION

Alterations of the gut microbiome influence the incidence and progression of not only gastric carcinogenesis (17), but also extra-intestinal cancers, such as breast and hepatocellular carcinoma, presumably through inflammatory and metabolic circuitries (18). Meanwhile, gut microbiota was also found contribute to the acute lung injury (19), the exacerbation of chronic obstructive pulmonary disease (COPD) (20), and the development of asthma (21), which highlighted its important role in affecting the respiratory system. Actually the hypothesis of “gut–lung” axis has been raised 20 years ago, when study found that gut-derived injurious factors can reach to the lung and systemic circulation via the intestinal lymphatics (22). Gut flora was found to be responsible for inducing lung inflammation against bacteria in mice and enhancing neutrophils infiltration through activation of tool like receptor 4 (TLR4) (23). Such immune transmission from gut to lung has been proved by many studies. One viewpoint supports that exacerbation of chronic lung diseases occur as an uncontrolled and inappropriate inflammatory response to bacteria colonizing damaged airways due to an ineffective Peyer’s patch-derived T lymphocyte response (24). These studies above strongly suggested a possible correlation between intestinal dysbiosis and the development of lung cancer. However, to our knowledge, no study has explored this possibility yet.

Although based on a small number of cases, we found a statistically significant decrease of alpha diversity in gut microbiota of lung cancer patients, compared with that of the healthy controls, which was consist with previous discovery that non-malignant lung tissues have higher microbiota alpha diversity than the paired tumors (10). The decline in both the bacterial diversity and richness was found in gut of patients with chronic inflammation such as inflammatory bowel disease (IBD) and colonrectal cancer, especially in patients with conventional adenoma (25). Reduced respiratory microbiome was also shown associated with greater emphysema and increased immune cell infiltration in COPD patients (26). These observations seem in line with the hygiene hypothesis that diverse microbes play an essential role in establishing the immune networks of a host, while in patients with various non-communicable inflammatory diseases, such as asthma, these regulatory networks seemed to be underrepresented and poorly developed (27).

Alterations of the gut microbial structure in lung cancer patients were characterized by the significant decrease of *Bacteroidetes* (phylum), *Bacteroidia* (Class), *Bacteroidales* (Order), and increase of *Firmicutes* (phylum), *Bacilli* (Class), *Actinomycetales*, and *Bacillales* (Order), which were mainly contributed by the reduction of bacteria genera such as *Prevotella* and *Dialister*, and the increase in *Ruminococcus, Streptococcus, Rothia, Bacillus, Actinomyces, Peptostreptococcus, etc.* These changes are not consistent with previous studies in sliva, bronchoalveolar lavage fluid, or lung tissues in patients with lung cancer, except that the genus of *Streptococcus* was previously found increased in lower airway of LC patients (28). But interestingly, they are highly agreed with the observations in patients with colon cancer that the class of *Bacilli*, genera of *Streptococcus, Actinomyces, Peptostreptococcus*, and etc are enriched (29). Importantly, we found a significantly increased ratio of *Bacteroides*/*Prevotella*, which was also proved in patients with colorectal cancer when compared with normal individuals (25). Taken together, these results indicate a state of dysbiosis in the gut microbiome of patients with lung cancer.

As the major genus that was found significantly reduced in LC patients’ gut, *Prevotella* had drawn many of our attentions. *Prevotella* strains are classically considered commensal bacteria due to the extensive presence in the healthy human body and the rare involvement in infections. *Prevotella* can stimulate epithelial cells to produce cytokines such as IL-8, IL-6 and CCL20, which can promote mucosal Th17 immune responses and neutrophil recruitment (30). Increased *Prevotella* abundance is associated with augmented Th17-mediated mucosal inflammation, which is in line with the marked capacity of *Prevotella* in driving Th17 immune responses through activation of TLR2 (30). These studies suggested an immune-stimulating activity of this genus. Congruously, *Prevotella* abundance was found reduced within the lung microbiota of patients with asthma and COPD (31), which further highlighted its important role in affecting lung cancer development.

On the other side, the increasing of certain bacteria in LC patients suggested a deleterious role of them in the development of cancer. Species of *Streptococcus* are found to increase in patients with LC. Compared to healthy controls, the NSCLC patients presented significantly higher frequencies of Th1 and Th17 cells reacting to *S. salivarius* and *S. agalactiae*, in the PB, LC, and GI tract (32). The order of *Actinomycetales* had been as potential colorectal cancer driver bacteria (25). Among them, the filamentous Gram-positive anaerobic genus *Actinomyces*, was found to cause *Actinomycosis*, which is a rare and slowly progressive infectious disease that can affect a variety of organ systems including the lung (33). *Actinomyces* was also found significantly associated with the carcinoma-in-adenoma group (34). Besides, *Peptostreptococcus* species, *P. stomatis* was confirming known to be associated with CRC (35). *P. anaerobius*, which is increased in human colon tumors compared with nontumor tissues, can enhance AOM-induced tumorigenesis in mice by activating TLR2/4-ROS-cholesterol axis (36). In addition, species of *Atopobium, A. parvalum* was found positively correlates with pediatric IBD disease severity (37). However, the effects of these bacteria on lung cancer development are still unclear, which deserves in-deep study.

In addition, we compared the gut microbiome of lung cancer patients according to different histological types based on collected samples, including SSC, ADC, and SCLC. Although the alpha diversity among groups showed no difference, we detected an obvious separation of beta diversity between patients with ADC and SCLC. Given the limited number of study cases, further large scale studies on the characterization of gut microbiome in LC patients with different histological types are necessary.

### Conclusions

In conclusion, this is the first report to show the alteration of gut microbiome in lung cancer patients. Our results showed that the gut microbiome of lung cancer patients altered significantly compared with healthy individuals. However, the association between microbial dysbiosis and lung cancer is not clearly understood, future studies involving larger cohorts and metagenomics, or metabolomics, may elucidate the correlations between gut microbiota and lung cancer development.

## MATERIALS AND METHODS

### Study subjects and sample collection

The recruitment of participants and the process of sample collection are depicted in Fig. 1. Fifty patients (age, 50–75 years) were ultimately recruited from the Second Affiliated Hospital of Dalian Medical University, Dalian, China, from September 2015 to July 2016. Fecal samples were collected in Stool Collection Tubes, which were pre-filled with Stool DNA Stabilizer for collection (Stratec, Germany), then frozen and stored at −80 °C for further use. All subjects were examined clinically before sampling and were subsequently divided into three groups: SCLC (n=17), AC (n=18), SCC (n=15). The samples of the healthy controls (HC, n = 30) were collected during routine physical examination at the First Affiliated Hospital of Dalian Medical University, Dalian, China.

The participants with the following diseases were excluded: cardiovascular disease, diabetes mellitus, liver cirrhosis, irritable bowel syndrome, inflammatory bowel disease, infections with known active bacteria, fungi, or virus. Those who abused drug or alcohol in the last year, or used antibiotics, probiotics, prebiotics, or synbiotics in the month, or received chemotherapy before collection of the fecal sample were also excluded.

### DNA extraction, polymerase chain reaction (PCR) and pyrosequencing

The microbial genome was extracted using E.Z.N.A. ® Stool DNA kit (Omega Bio-tek, Inc.) according to the manufacturer’s instructions. A Nanodrop 2000 Spectrophotometer was used to evaluate the purity and concentration of isolated DNA. The polymerase chain reaction (PCR) to amplify the V3-V4 region of bacterial 16S ribosomal RNA gene was performed as described previously. After amplicons extraction, samples were purified using AXYGEN gel extraction kit (Qiagen) and quantified by Quant-iT PicoGreen dsDNA Assay Kit on Microplate reader (BioTek, FLx800). Sequencing and data analysis were subsequently performed on an Illumina MiSeq platform by Personal Biotechnology Co., Ltd. (Shanghai, China). The taxa classification and statistical analysis were conducted as described in previous studies.

### Statistics and analysis

Illumina MiSeq sequences obtained after quality control analysis were used in the present analysis, which were uploaded to QIIME (Quantitative Insights Into Microbial Ecology, v1.8.0) for further study. The operational taxonomy units (OTUs) of representative sequences at a similarity cutoff of 97% and their relative abundance (α-diversity) were used to calculate ACE and Chao1 index by UCLUST. The abundance and diversity of the OTUs (β-diversity) were examined using principal component analysis (PCA), Principal coordinates analysis (PCoA) and nonmetric multidimensional scaling (NMDS) with weighted and unweighted UniFrac analysis in R software. The statistical significance of the separation among groups was assessed by the linear discriminant analysis effect size (LEfSe) method based on linear discriminant analysis scores exploited by Curtis Huttenhower (http://huttenhower.sph.harvard.edu/galaxy/), which used the nonparametric factorial Kruskal–Wallis and Wilcoxon rank sum test to identify key OTUs for separating different treatment groups at a significance level of 0.05. Metastats analysis was performed based on the raw count data matrix to find out the taxa statistically different between HC and LC samples.

For the analyses of clinical data, the non-parametric *t*-test between HC and LC groups was performed with the assistance of GraphPad Prism 6 (Graph Pad Software, La Jolla, CA, USA). Results were considered to be statistically significant with P < 0.05. The taxa classification and statistical analysis were conducted as described in previous studies (38).

### Ethics statement

This study protocol was approved by the Ethics Committee of the Second Affiliated Hospital of Dalian Medical University, Dalian, China. After receiving a written description of the aim of this study, all participants gave written informed consent prior to enrollment.

## SUPPLEMENTARY MATERIAL

### Supplementary Figure legends

**FIG S1**, Comparison of the Shannon **(A)** and Simpson **(B)** index of gut microbiome in HC and LC groups.

**FIG S2**, The gut bacterial composition in different groups at Phylum, Class, Order, and Family levels.

**FIG S3**, Principal component analysis of gut microbiome in HC and LC groups. PC1 and PC2 account for 73.08% of the variation.

**FIG S4**, Hierarchical clustering based on Weighted UniFrac by the method of UPGMA. Number of samples: HC, 2-20, LC, 62-98

**FIG S5**, The relative abundance of the most differentially abundant bacterial taxons identified by LEfSe in each samples of different among groups.

**FIG S6**, The specific bacterial Phyla, Classes, and Orders that are significantly changed in LC patients compared with HCs detected by the MetaStat method.

**FIG S7**, Alpha diversity of gut microbiota in LC patients with specific histological types. **(A)** Venn diagram of shared and independent bacterial OTUs in different groups. **(B)** ACE, **(C)** Chao1, **(D)** Shannon, **(E)** Simpson.

**FIG S8**, The taxonomy-based comparison among groups at the genus level showed by heat map.

#### ACKNOWLEDGMENTS

This study was supported by the National Natural Science Foundation of China (NSFC, 81370113), the Nature Science Foundation of Liaoning Province, China (2015020262), and the Research Foundation from the Department of Education, Liaoning Province, China (L2016003). This work was supported by Liaoning Provincial Program for Top Discipline of Basic Medical Sciences.

## CONFLICTS OF INTERSTS

The authors declare no competing interest.

## REFERENCES

1. Herbst RS, Heymach JV, Lippman SM. 2008. Lung cancer. N Engl J Med 359(13): 1367–1380.

2. Siegel RL, Miller KD, Jemal A. Cancer statistics, 2016. 2016. CA Cancer J Clin 66(1): 7–30.

3. Youlden DR, Cramb SM, Baade PD. 2008. The international epidemiology of lung cancer: geographical distribution and secular trends. J Thorac Oncol 3(8): 819–831.

4. Wang P, Zou J, Wu J, Zhang C, Ma C, Yu J, Yu J, Zhou Y, Li B, Wang K. 2017. Clinical profiles and trend analysis of newly diagnosed lung cancer in a tertiary care hospital of East China during 2011-2015. J Thorac Dis 9(7): 1973–1979.

5. Wang X, Pittman GS, Bandele OJ, Bischof JJ, Liu G, Brothers JF 2nd, Spira A, Bell DA. 2017. Linking polymorphic p53 response elements with gene expression in airway epithelial cells of smokers and cancer risk. Hum Genet 133(12): 1467–1476.

6. Man WH, de Steenhuijsen Piters WA, Bogaert D. 2017. The microbiota of the respiratory tract: gatekeeper to respiratory health. Nat Rev Microbiol 15(5): 259–270.

7. Roudi R, Mohammadi SR, Roudbary M, Mohsenzadegan M. 2017. Lung cancer and beta-glucans: review of potential therapeutic applications. Invest New Drugs 35(4): 509–517.

8. Lee SH, Sung JY, Yong D, Chun J, Kim SY, Song JH, Chung KS, Kim EY, Jung JY, Kang YA, Kim YS, Kim SK, Chang J, Park MS. 2016. Characterization of microbiome in bronchoalveolar lavage fluid of patients with lung cancer comparing with benign mass like lesions. Lung Cancer 102: 89–95.

9. Yu G, Gail MH, Consonni D, Carugno M, Humphrys M, Pesatori AC, Caporaso NE, Goedert JJ, Ravel J, Landi MT. 2016. Characterizing human lung tissue microbiota and its relationship to epidemiological and clinical features. Genome Biol 17(1): 163.

10. Chen MM, Zahs A, Brown MM, Ramirez L, Turner JR, Choudhry MA, Kovacs EJ. 2014. An alteration of the gut-liver axis drives pulmonary inflammation after intoxication and burn injury in mice. Am J Physiol Gastrointest Liver Physiol 307(7): G711–718.

11. Trompette A, Gollwitzer ES, Yadava K, Sichelstiel AK, Sprenger N, Ngom-Bru C, Blanchard C, Junt T, Nicod LP, Harris NL, Marsland BJ. 2014. Gut microbiota metabolism of dietary fiber influences allergic airway disease and hematopoiesis. Nat Med 20(2): 159–166.

12. Zou Y, Dong C, Yuan M, Gao G, Wang S, Liu X, Han H, Li B. 2014. Instilled air promotes lipopolysaccharide-induced acute lung injury. Exp Ther Med 7(4): 816–820.

13. Ben DF, Yu XY, Ji GY, Zheng DY, Lv KY, Ma B, Xia ZF. 2012. TLR4 mediates lung injury and inflammation in intestinal ischemia-reperfusion. J Surg Res 174(2): 326–333.

14. Kim YG, Udayanga KG, Totsuka N, Weinberg JB, Núñez G, Shibuya A. 2014. Gut dysbiosis promotes M2 macrophage polarization and allergic airway inflammation via fungi-induced PGE(2). Cell Host Microbe 15(1): 95–102.

15. Sze MA, Tsuruta M, Yang SW, Oh Y, Man SF, Hogg JC, Sin DD. 2014. Changes in the bacterial microbiota in gut, blood, and lungs following acute LPS instillation into mice lungs. PLoS One 9(10): e111228.

16. He Y, Wen Q, Yao F, Xu D, Huang Y, Wang J. 2017. Gut-lung axis: The microbial contributions and clinical implications. Crit Rev Microbiol 43(1): 81–95.

17. Huang X, Li C, Li F, Zhao J, Wan X, Wang K. 2018. Cervicovaginal microbiota composition correlates with the acquisition of high-risk human papillomavirus types. Int J Cancer 143(3):621–634.

18. Coker OO, Dai Z, Nie Y, Zhao G, Cao L, Nakatsu G, Wu WK, Wong SH, Chen Z, Sung JJY, Yu J. 2017. Mucosal microbiome dysbiosis in gastric carcinogenesis. Gut 67(6):1024–1032.

19. Mima K, Nakagawa S, Sawayama H, Ishimoto T, Imai K, Iwatsuki M, Hashimoto D, Baba Y, Yamashita YI, Yoshida N, Chikamoto A, Baba H. 2017. The microbiome and hepatobiliary-pancreatic cancers. Cancer Lett 402: 9–15.

20. Nicod LP, Kolls JK. 2015. Chair’s summary: mechanisms of exacerbation of lung diseases. Ann Am Thorac Soc 12 Suppl 2: S112–114.

21. Ottiger M, Nickler M, Steuer C, Bernasconi L, Huber A, Christ-Crain M, Henzen C, Hoess C, Thomann R, Zimmerli W, Mueller B, Schuetz P. 2017. Gut, microbiota-dependent trimethylamine-N-oxide is associated with long-term all-cause mortality in patients with exacerbated chronic obstructive pulmonary disease. Nutrition 45:135–141.e1.

22. Kang YB, Cai Y, Zhang H. 2017. Gut microbiota and allergy/asthma: From pathogenesis to new therapeutic strategies. Allergol Immunopathol (Madr) 45(3): 305–309.

23. Magnotti LJ, Upperman JS, Xu DZ, Lu Q, Deitch EA. 1998. Gut-derived mesenteric lymph but not portal blood increases endothelial cell permeability and promotes lung injury after hemorrhagic shock. Ann Surg 228(4): 518–527.

24. Tsay TB, Yang MC, Chen PH, Hsu CM, Chen LW. 2011. Gut flora enhance bacterial clearance in lung through toll-like receptors 4. J Biomed Sci 18: 68.

25. Samuelson DR, Welsh DA, Shellito JE. 2015. Regulation of lung immunity and host defense by the intestinal microbiota. Front Microbiol 6: 1085.

26. Peters BA, Dominianni C, Shapiro JA, Church TR, Wu J, Miller G, Yuen E, Freiman H, Lustbader I, Salik J, Friedlander C, Hayes RB, Ahn J. 2016. The gut microbiota in conventional and serrated precursors of colorectal cancer. Microbiome 4(1): 69.

27. Richmond BW, Brucker RM, Han W, Du RH, Zhang Y, Cheng DS, Gleaves L, Abdolrasulnia R, Polosukhina D, Clark PE, Bordenstein SR, Blackwell TS, Polosukhin VV. 2016. Airway bacteria drive a progressive COPD-like phenotype in mice with polymeric immunoglobulin receptor deficiency. Nat Commun 7: 11240.

28. Smits HH, Hiemstra PS, Prazeres da Costa C, Ege M, Edwards M, Garn H, Howarth PH, Jartti T, de Jong EC, Maizels RM, Marsland BJ, McSorley HJ, Müller A, Pfefferle PI, Savelkoul H, Schwarze J, Unger WW, von Mutius E, Yazdanbakhsh M, Taube C. 2016. Microbes and asthma: Opportunities for intervention. J Allergy Clin Immunol 137(3): 690–697.

29. Liu HX, Tao LL, Zhang J, Zhu YG, Zheng Y, Liu D, Zhou M, Ke H, Shi MM, Qu JM. 2018. Difference of lower airway microbiome in bilateral protected specimen brush between lung cancer patients with unilateral lobar masses and control subjects. Int J Cancer 142(4):769–778.

30. Sobhani I, Tap J, Roudot-Thoraval F, Roperch JP, Letulle S, Langella P, Corthier G, Tran Van Nhieu J, Furet JP. 2011. Microbial dysbiosis in colorectal cancer (CRC) patients. PLoS One 6(1): e16393.

31. Larsen JM. 2017. The immune response to Prevotella bacteria in chronic inflammatory disease. Immunology 151(4): 363–374.

32. Hilty M, Burke C, Pedro H, Cardenas P, Bush A, Bossley C, Davies J, Ervine A, Poulter L, Pachter L, Moffatt MF, Cookson WO. 2010. Disordered microbial communities in asthmatic airways. PLoS One 5(1): e8578.

33. Ma QY, Huang DY, Zhang HJ, Wang S, Chen XF. 2017. Upregulation of bacterial-specific Th1 and Th17 responses that are enriched in CXCR5 (+) CD4 (+) T cells in non-small cell lung cancer. Int Immunopharmacol 52: 305–309.

34. Laguna S, Lopez I, Zabaleta J, Aguinagalde B. 2017. Actinofmycosis associated with foreign body simulating lung cancer. Arch Bronconeumol 53(5): 284–285.

35. Kasai C, Sugimoto K, Moritani I, Tanaka J, Oya Y, Inoue H, Tameda M, Shiraki K, Ito M, Takei Y, Takase K. 2016. Comparison of human gut microbiota in control subjects and patients with colorectal carcinoma in adenoma: Terminal restriction fragment length polymorphism and next-generation sequencing analyses. Oncol Rep 35(1): 325–333.

36. Yu J, Feng Q, Wong SH, Zhang D, Liang QY, Qin Y, Tang L, Zhao H, Stenvang J, Li Y, Wang X, Xu X, Chen N, Wu WK, Al-Aama J, Nielsen HJ, Kiilerich P, Jensen BA, Yau TO, Lan Z, Jia H, Li J, Xiao L, Lam TY, Ng SC, Cheng AS, Wong VW, Chan FK, Xu X, Yang H, Madsen L, Datz C, Tilg H, Wang J, Brünner N, Kristiansen K, Arumugam M, Sung JJ, Wang J. 2017. Metagenomic analysis of faecal microbiome as a tool towards targeted non-invasive biomarkers for colorectal cancer. Gut 66(1): 70–78.

37. Tsoi H, Chu ESH, Zhang X, Sheng J, Nakatsu G, Ng SC, Chan AWH, Chan FKL, Sung JJY, Yu J. 2017. Peptostreptococcus anaerobius induces intracellular cholesterol biosynthesis in colon cells to iduce proliferation and causes dysplasia in mice. Gastroenterology 152(6): 1419–33 e5.

38. Mottawea W, Chiang CK, Mühlbauer M, Starr AE, Butcher J, Abujamel T, Deeke SA, Brandel A, Zhou H, Shokralla S, Hajibabaei M, Singleton R, Benchimol EI, Jobin C, Mack DR, Figeys D, Stintzi A. 2016. Altered 31.intestinal microbiota-host mitochondria crosstalk in new onset Crohn’s disease. Nat Commun 7: 13419.

